# Multipurpose virtual reality environment for biomedical and health applications

**DOI:** 10.1101/366302

**Authors:** Jordi Torner, Stavros Skouras, José L. Molinuevo, Juan D. Gispert, Francisco Alpiste

## Abstract

Virtual reality is a trending, widely accessible, contemporary technology of increasing utility to biomedical and health applications. However, most implementations of VR environments are tailored to specific applications. We describe the complete development of a novel, open-source virtual reality environment that is suitable for multipurpose biomedical and healthcare applications. The developed environment simulates an immersive (first-person perspective) run in the countryside, in a virtual landscape with various salient features. The utility of the developed VR environment has been validated via two test usage cases: an application in the context of motor rehabilitation following injury of the lower limbs and an application in the context of real-time functional magnetic resonance imaging neurofeedback, to regulate brain function in specific regions of interest. The resulting test applications suggest that the implemented approach is robust, versatile and efficient. Both applications are publicly available via a GitHub repository, in support of the Open Science initiative. We anticipate our contribution to catalyze further progress and replicability with regards to the usage of virtual reality in biomedical and health applications.

*Index Terms*— Motor rehabilitation, neurofeedback, virtual reality.

## I. Introduction

Virtual reality (VR) environments have been in use in training applications, for over four decades. In recent times, the active development of 3D technologies concerned with medical therapy, training and rehabilitation has become increasingly important as an area of interest in research and healthcare. Moreover, biomedical applications using the VR technology are receiving increasing attention and are becoming increasingly accessible to consumers [1-2]. User studies show that such applications are both effective and intuitive [3-4]. The development of VR has followed a very interesting technological history reliant on a series of independent engineering innovations.

In 1844 Sir C. Wheatstone invented the “stereoscope” [5]. By obtaining two identical photographs from different angles and presenting the leftmost image to the left eye and the rightmost image to the right eye, a 3D perceptual effect was created. In 1891, L. D. du Hauron patented the “anaglyph” [6], which consisted of eliminating the red color for the images of the right eye and eliminating the green and blue colors for those of the left eye. In this way a 3D percept was effectively introduced to the images. In 1961, C. P. Corneau and J. S. Bryan, built the first VR helmet [7]. The device was equipped with a magnetic sensor that determined the orientation of the user’s head and allowed viewing images stereoscopically, in reference to the direction of gaze. Nevertheless, the origin of VR is usually attributed to 1965, when I. E. Sutherland published an article entitled “The Ultimate Display” in which he elaborately described the concept of VR [8]. VR was further enhanced with the advent of computer graphics, largely thanks to the work of L. G. Roberts and I. E. Sutherland in the 60s and 70s [9-10]. The context in which VR found its first industrial application was that of flight simulators. In 1981 T. Furness developed the “Virtual Cabin” [11]. It was the first simulator of the airplane cabin used to train pilots. From the 80s onwards the mass-scale availability of VR technology has been increasing steadily.

At present, VR is a technology that uses smartphones or specialized headsets in combination with physical spaces or multi-projected environments, to generate realistic images, sounds and sensations that simulate a user’s physical presence in a virtual stage. It offers users the possibility to experience psychological states of immersion and involvement and gives rise to a sense of presence. In the most advanced contemporary VR applications, the user experience is enhanced by the technological capacity to map physical movement into the VR environment through the use of accelerometers and other movement-sensing devices, as well as by the addition of other innovative features that complement a multi-sensory immersive experience (e.g. stimulation via temperature changes, olfactory information, manipulation of proprioceptive state, neurofeedback, etc.).

## II. Related Work

Due to the relative accessibility of VR and due to its ability to provide an experience with high ecological validity, VR has been used in a variety of biomedical and healthcare applications. Such applications range from psychiatric treatments to rehabilitation and from medical education to surgical training. Notable examples of psychiatric applications include the treatment of phobias [12], the training of emotion regulation [13] and the training of autistic social skills [14]. Additionally, immersive technologies have been used to support cognitive training in older adults with mild cognitive impairment or dementia [15]. VR training can be effective in improving global cognition, specific cognitive domains and psychosocial functioning in people with mild cognitive impairment [16] and dementia [17], including dementia due to Alzheimer’s disease [18]. VR games in which the goal is accomplished by activating specific muscles, thereby facilitating rehabilitation physiotherapy [19-20], comprise another good example of the rapidly expanding field of “serious gaming” using VR [21-22].

In the field of surgery, in cases of complex operations, tools have been developed to maximize the probability of a positive outcome. VR applications can enable better vision and increased control [23]. Simulations of surgical procedures have been used in training for laparoscopic, robotic, gynecological, neurological and cardiac surgery [24]. In medical education, information on a disease can be directed to different groups (doctors, students or families) and learning is facilitated via first-person perspectives in immersive, realistic environments [25]. Among numerous other applications, VR has been utilized successfully in the rehabilitation of patients with stroke [26–28]. Neurorehabilitative therapies that focus on motor recovery have utilized electroencephalography and brain-computer interfaces featuring 3D virtual avatars for controlling simulated movement [29], leading into the field of Virtual Reality Control [30].

Despite the merits of VR and its extensive utility, the vast majority of previously implemented VR environments are tailored to one application or situation and are not designed in a way that allows for easy modification and adaptation to new settings. We have developed a new, open-source environment that is suitable for multipurpose biomedical and health applications. This VR environment emulates an immersive (first-person perspective) run in the countryside, in a graphical environment with various salient features (e.g. a lake, a bridge, trees and mountains, etc.; Fig. 1). Two user cases have been tested: a first test application in the context of motor rehabilitation following injury and another in the context of neurofeedback with real-time magnetic functional imaging. Starting with these particular applications was motivated by previous reports of successful VR-facilitated motor rehabilitation [4, 19] and successful self-regulation of brain function using VR to represent changes in brain activity [31-32].

**Fig. 1.**
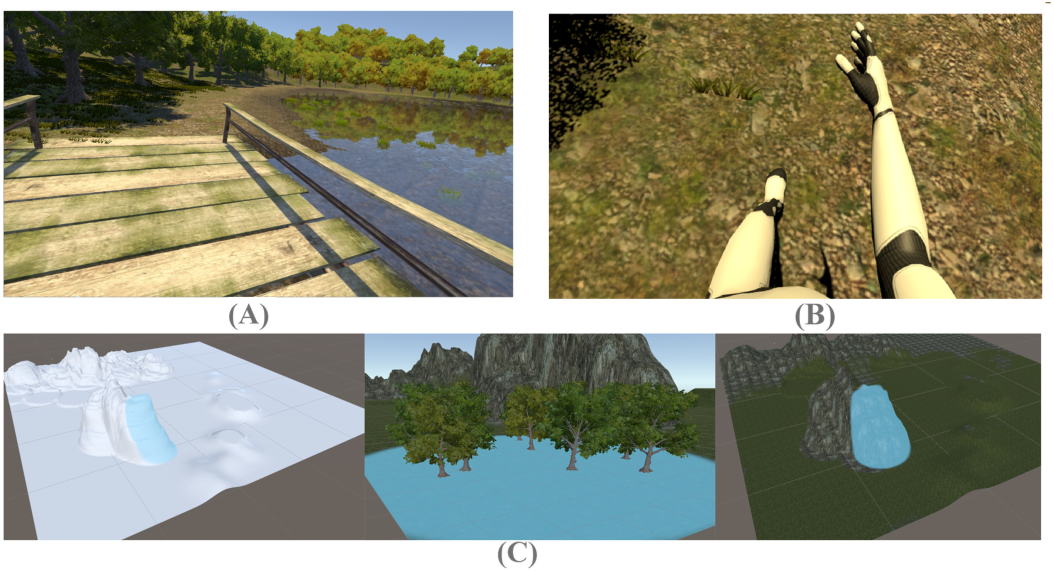
The designed VR environment. An instance of the landscape typically seen by participants (**A**); illustration of the immersive, first-person perspective (**B**); three snapshots from the graphics design process (**C**).

## III. Methods

In this section we present the design of the 3D computer graphics and the data management options for the multipurpose VR environment. Subsequently, we present the motivation, apparatus, procedure and computations associated to each of the first two test applications.

The multipurpose VR environment has been entirely created using Unity 5.5.0 [33]. Unity is a game engine that is widely used for video game applications on multiple platforms, from PCs and gaming consoles to mobile devices and websites. With Unity, it is possible to create 2D or 3D applications using object-oriented programming, via the C, C++ and C Sharp programming languages, and each object provides a specific functionality. The following procedure was used for the design of the computer graphics. Starting with a completely flat terrain, the first step was to use the modeling tools to give shape to the landscape. The relief was modified by adding textures, trees and grass through the use of specialized modeling Unity tools. An avatar character was created using the Autodesk Character Generator [34]. The possibility of using the application in first-person view or third-person mode was enabled by the inclusion of several different virtual cameras (Fig.1a).

An application using the developed multipurpose VR environment, can optionally include a graphical user interface (GUI) for data management. Via the data management GUI, data protection via username/password login can be enabled, a feature that was developed with applications related to confidential medical data in mind. A new study can be selected for new patients or a new session for returning patients. Stored data from previous visits are easily accessible via a specialized “Studies” tab. A “patient management” tab can be used to add new patients and to access a patient’s performance history. Another optional feature, regards a server database application developed in Hypertext Preprocessor scripting language (PHP), that can collect information about the institution where the software is used as well as the performance statistics, facilitating multi-center collaborative research and/or clinical monitoring in multiple locations.

### A. Test application 1: Motor rehabilitation for lower limbs

The aim of the first test application developed using the multipurpose VR environment is to make the process of rehabilitation more enjoyable and effective, for patients who have suffered an injury or who have another mobility problem involving their leg(s). Subjects can be immersed in the VR environment from a first-person visual perspective, so that they have the sensation that the virtual legs are their own. Using four accelerometers placed on a subject’s physical legs, the application can detect movement and apply it to the virtual legs, resulting in movement within the VR environment.

The software application has been created using Unity 5.5, as aforementioned and works with four accelerometers connected via bluetooth and controlled by an Arduino board. During the software development, we used the Oculus Rift DK2 VR glasses, featuring a resolution of 2160 × 1200 pixels and a vision angle of 110° [35]. Additionally, an Intel Core i7-5300 computer workstation with 8GB RAM, 1TB storage, and a NVIDIA GeForce GTX 970 high performance graphics card was used. The developed software source code can be easily adapted for usage with other devices. The minimum computer requirements comprise of an Intel5 processor, 8GB of RAM and a NVIDIA GeForce GTX 960 or similar high performance graphics card. Moreover, the required apparatus must incorporate an accelerometer and a gyroscope that can read data in three reference axes, similarly to the detailed connection scheme displayed in Fig. 2. A calibration of the accelerometers is necessary for each patient during the initiation of a session. During the software development, using the Arduino Pro Mini ATmega 328P was preferred due to its compact dimensions of 18 × 33 mm. This model operates at 5V and 16 MHz. For the purposes of the laboratory setup, pins were welded in-house on a custom board with 14 pins of input/output, 6 analog inputs and additional space for mounting other connectors. The hardware configuration consisted of four Inertial Measurement Unit (IMU) devices. IMUs are electronic devices that measure and report the speed, orientation and gravitational forces of a device, using an accelerometer and a gyroscope. Specifically, the MPU-6050 (Invensense Inc.) model, used during development, has six degrees of freedom. It combines a three-axis accelerometer and a three-axis gyroscope and operates at a sampling frequency of 100 Hz using 3.3V. The Bluetooth module JY-MCU HC-06, that is fully compatible with the Arduino, was selected due to its compact size that deemed it ideal for the project. Finally, a FDPI USB-to-TTL serial cable, with an embedded FTDI FT232RL USB-serial chip, featuring a six-pin socket with 5V power and ground, as well as RX, TX, RTS and CTS connectors at 3V logic levels, completed the VR lab setup (Fig. 2).

**Fig. 2.**
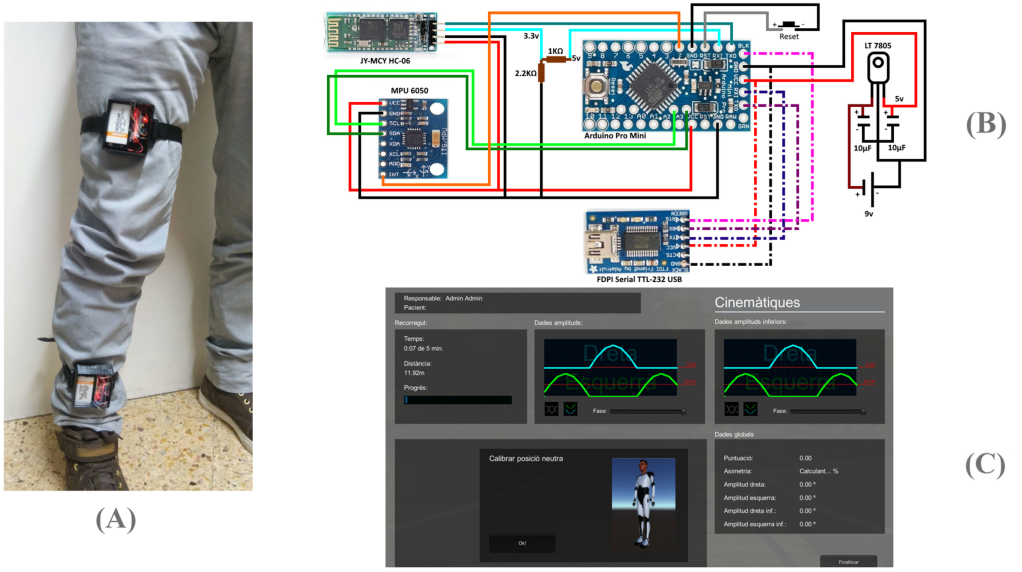
Application to motor rehabilitation. Device mounting (**A**); hardware connection scheme (**B**); real-time data (**C**).

During a VR motor rehabilitation session, a patient can be immersed in the landscape in the form of an avatar, in first-person perspective, so that the patient has the sensation that the virtual legs are his/her own (Fig 1). Using four accelerometers placed on the patient’s legs (Fig. 2), the application can detect movement and map the movement to the VR environment. Regardless of the magnitude of the movement of the patient’s leg, i.e. anywhere between a twitch and a complete step, each leg movement is mapped into a complete step in the VR environment. This way, patients with limited mobility can have the rewarding and encouraging experience of walking and jogging in a virtual environment. Each round of practice should typically last for a maximum of five minutes.

The data from a session using the VR environment for rehabilitation following leg injury are updated in real-time, as well as stored and summarized on a screen at the end of each session. The information displayed includes: a) The patient name or code and the name of the researcher or clinician; b) Real-time updates of the distance travelled, the time taken, and the remaining length of the session; c) Real-time information on the current amplitudes of the superior (above the knee) and inferior (below the knee) angle of flexion of each leg; d) Real-time information on the average amplitudes of the superior and inferior angle of flexion of each leg, since the beginning of the session; e) Real-time information on the asymmetry α, in accordance with (1) and (2).

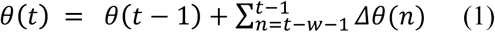

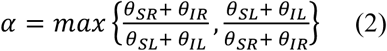

where *θ(t)* represents the angular position in a measurement location at time *t, Δθ* represents the change in angular position between successive sampling points, *w* represents the angular position update window and SR, IR, SL, IL represent the measurement locations Superior-Right, Inferior-Right, Superior-Left and Inferior-Left, respectively. For the purposes of the application, *w* was set to 10 samples; that is the angular velocity was updated every one second.

### B. Test application 2: Real-time functional neurofeedback

Motivated by recent reports of successful self-regulation of brain activity using VR environments, particularly with electroencephalography [31] and functional near-infrared spectroscopy [32], we developed a VR real-time functional magnetic resonance imaging neurofeedback paradigm (rt-fMRI NF). The speed of movement in the VR environment was modulated by the efficiency of self-regulation of brain activity. The specified brain region of interest (ROI) was the CA1 subfield of the hippocampus, an area that is expected to show changes in brain function during the early, asymptomatic stage of Alzheimer’s disease [36]. The local ethics committee “CEIC-Parc de Salut Mar” reviewed and approved the experimental protocol used during a pilot trial.

A Philips Ingenia CX MRI scanner, operating at 3 Tesla, was used, featuring Philips console software version 5.1 and the Philips external control (XTC) patch. Apart from the Philips image reconstruction and MRI console PCs, two additional computers (Intel7 processors, 16GB RAM), running Windows 7 Enterprise edition and in-house neuroimaging analysis software with dependencies on Matlab 2013b (Mathworks Inc.) and the SPM12 toolbox [37], were additionally used. The PCs were connected as illustrated in Fig. 4. Apart from the image reconstruction PC, the rest of the computers were connected via shared folders through the local area network (LAN) and file transfer was facilitated through MS-DOS batch programs and the Philips “Corba Data Dumper” application. The digital audio-visual stimulation system VisuaStimDigital (Resonance Technology Inc.), featuring MRI-compatible goggles with fiber-optic communication, were used to present the VR environment to the eyes of a volunteer undergoing functional MRI scanning.

Scanning was performed on the 3 T Philips Ingenia CX with a 32-channel headcoil. Prior to the functional MRI measurements, a high-resolution (1×1×1 mm) T1-weighted anatomical reference image was acquired using a rapid acquisition gradient echo sequence. Echo planar imaging was used with an echo time of 35 ms and a repetition time (TR) of 3 s. Slice-acquisition was interleaved within the TR interval. The matrix acquired was 80×80 voxels with a field of view of 240 mm, resulting in an in-plane resolution of 3 mm. Slice thickness was 3 mm with an interslice gap of 0.2 mm (45 slices, whole brain coverage).

Each EPI image of whole-brain activity was acquired, reconstructed and exported with minimal delay, every three seconds, via the local area network (LAN), to the real-time analysis computer (Fig. 4). Movement correction through rigid-body registration to an initial reference volume, temporal high-pass filtering with a cutoff frequency of 1/200 Hz to remove low frequency drifts in the fMRI time series [38], and voxel efficiency weighting [39] through voxel-wise normalization within a sliding time-window, were applied in real-time to the online data. Voxel-efficiency weighting was performed by normalizing new images based on the mean and standard deviation of the preceding observations in each voxel, according to (3).

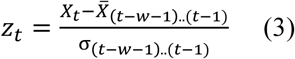

where, at time *t, z_t_* represents the efficiency-weighted value in a single voxel, *X_t_* represents the unweighted value of that voxel, 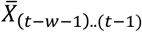 represents the mean and *σ*_(_*_t_*_−_*_w_*_−1).(_*_t_*_−1)_ represents the standard deviation of the timeseries within a specified time-window, of length *w*, up to time *(t* − 1). For the purposes of our testing, *w* was set to 30 scanning volumes.

An outcome variable was instantiated for the activity measured within a region of interest (ROI), for each acquired brain volume, according to the following computations. The expected data vector **Y** at time t_0_ was computed as in (4).

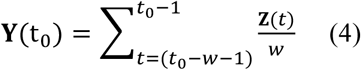

where **Z**(t) represents the voxel-efficiency-weighted observed data vector from the ROI at time t and *w* represents the length of the specified sliding time-window (30 volumes). For each new volume acquired in real-time, a non-linear metric *NF* was computed according to (5).

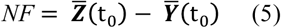

When the average expected signal in the ROI surpassed the average observed signal of the ROI in the new volume, the velocity *v* in the VR environment was increased by 5% as in (5); in the opposite case, the velocity was decreased by 5% as in (6).

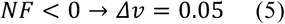

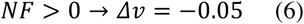

where *v* represents the velocity of movement in the VR environment.

This procedure resulted in a closed-loop, non-linear, adaptive, sliding window paradigm, enabling continuous neurofeedback throughout the scanning session. The first 10 scanning volumes were discarded to allow for magnetic field saturation. The initial baseline ROI mean and standard deviation were computed based on the following 30 functional volumes, during which the VR velocity was maintained at 50%. After the initial reference baseline was established, every three seconds, the real-time neuroimaging analysis variable was communicated to the VR presentation PC, resulting in a modulation of the velocity of movement in the VR environment (Fig. 4). The velocity of movement was increased by 5% when changes in the neural activity within the ROI were in the desired direction or decreased by 5% when changes in the neural activity within the ROI were in the opposite direction. The velocity would remain stable in the highly improbable case of no change in neural activity.

## IV. Results

In this section, we present the results available from the usage of each of the first two test applications that have been developed based on the multipurpose VR environment.

### A. Test application 1: Motor rehabilitation for lower limbs

During a VR motor rehabilitation session, real-time results are displayed in a screen that is only visible by the investigator or clinician, because the patient is immersed in the VR environment, viewing through the VR goggles. After the end of the VR session, the final results summarize the performance data and overall progress, in relation to previous sessions. Selecting a particular metric (e.g. “Puntuacio”, i.e. Score; Fig. 3) displays the history of performance on that metric across sessions. This setup allows to easily monitor progress and to intuitively identify the exact sources of improvement, as well as the muscles that require more exercise.

**Fig. 3.**
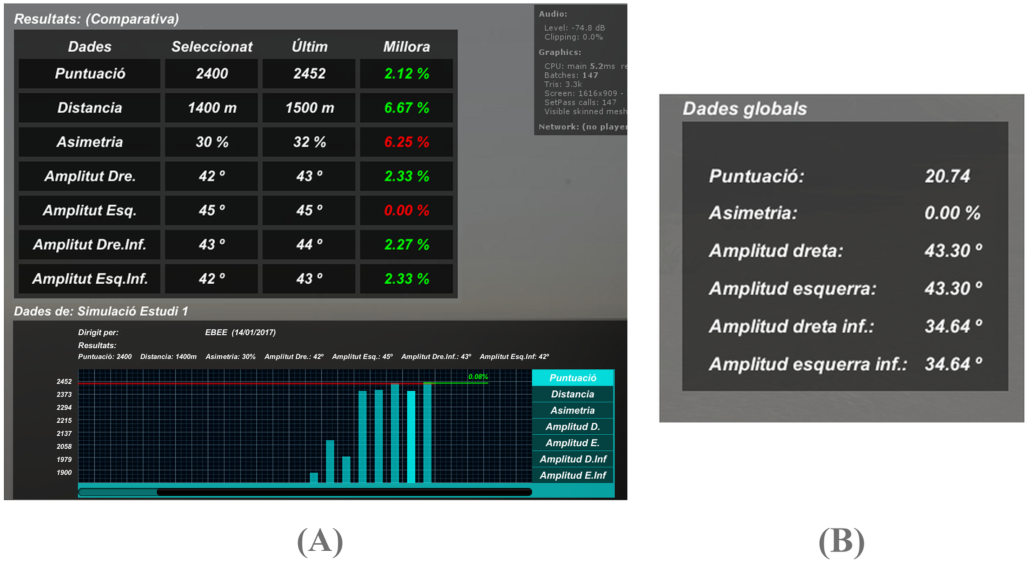
Exemplary results from a VR-facilitated motor rehabilitation session (**A**); global data computed (**B**).

### B. Test application 2: Real-time functional neurofeedback

During a VR real-time neurofeedback session, real-time results are displayed in a screen that is only visible by the investigator or clinician, because the subject is immersed in the VR environment, viewing through the VR goggles. Available real-time data include the three translation movement parameters in mm (X, Y, Z), the three rotation parameters in degrees (pitch, roll, yaw) and the non-linear metric *NF* that determines neurofeedback updates (Fig. 4), computed as in (5).

**Fig. 4.**
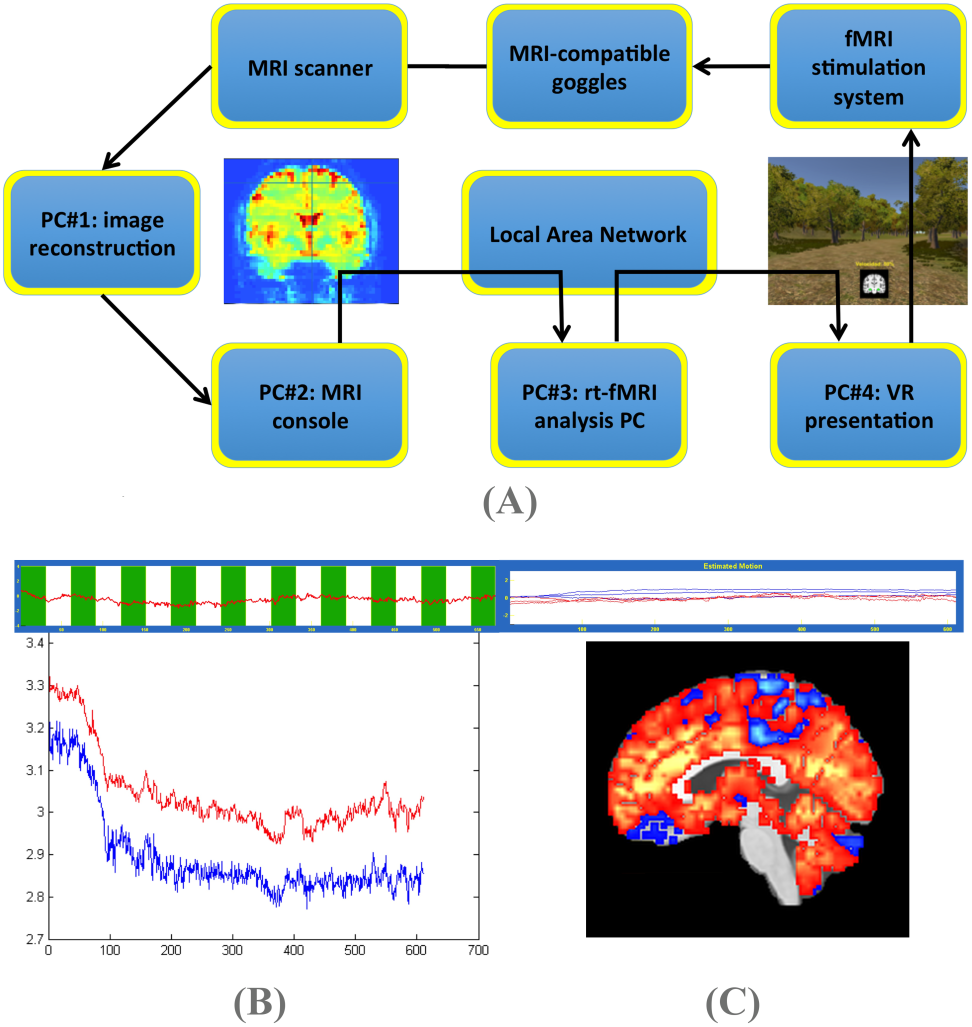
Application to real-time neuroimaging. In the real-time fMRI neurofeedback setup, the computers handling data reconstruction, data storage, data processing and presentation of the VR environment are interconnected serially or via the local area network (**A**). The real-time information available to investigators includes non-linear ROI signal change (top-left), motion parameters (top-right), ROI mean and median (bottom-left) (**B**). An innovative feature of the implemented paradigm is that it enables to investigate the whole-brain functional connectivity of the region of interest during or after the VR neurofeedback self-regulation of neural activity (**C**).

**Fig. 5.**
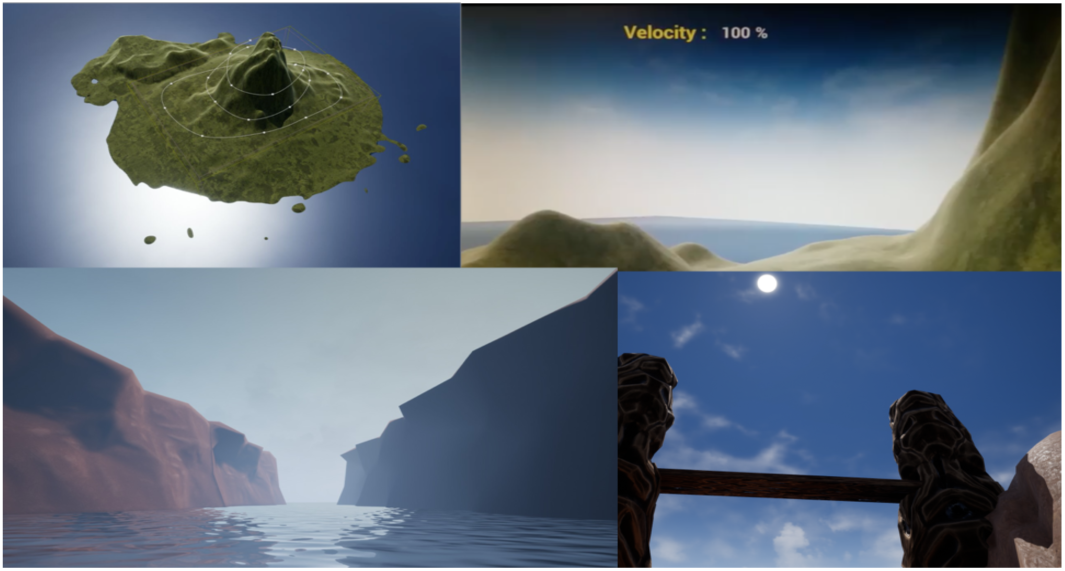
Future directions. Graphical snapshots from the upcoming, second generation multipurpose VR environment, featuring additional activities (i.e. swimming and flying), designed to enable versatile training within a single session.

At the end of the session, a VR score is immediately accessible, summarizing the distance travelled in the VR environment, which reflects neurofeedback performance. The neuroimaging data are also available for offline analyses to investigate the average activity within the ROI or the functional connectivity of the ROI with the rest of the brain (Fig. 4). Additionally, a log file is generated at the end of each session, documenting the sequential velocity values experienced by the subject.

## V. Discussion

To our knowledge, this is the first open-source, adaptive, multipurpose VR environment that can be used for several, different biomedical and health applications. The framework developed, provides features of clinical relevance, such as password protection options for data confidentiality, history logging and session management tools, automated performance tracking and real-time monitoring.

Due to the capacity VR offers for sensory immersion, the presented VR environment can be used for neurofeedback training as well as to facilitate rehabilitation following leg injury. In both cases, the user’s activity controls virtual movement. In the case of rehabilitation, the calibrated displacement of the patient is reflected through a virtual avatar with synchronous movement. Virtual steps are typically of greater magnitude than the triggering physical movements, thereby encouraging progress in recovery. In the case of rt-fMRI neurofeedback, changes in brain function processed in real-time, control the speed of movement in the VR environment, reinforcing learning of self-regulation with regards to the activity within a pre-specified brain area.

A minor limitation of the current version of the VR environment, is that it can become repetitive when used for extensive time periods. A variety of virtual landscapes and virtual objects are currently being designed to alleviate this identified shortcoming and to ensure a more fun and memorable experience, even during long sessions. The forthcoming, second edition of the presented VR environment, aims to encompass further innovative features, such as simultaneous representation of activity from multiple sources (e.g. upper and lower limbs or two independent brain regions).

In the context of motor rehabilitation, an application for rehabilitation following injury of the upper limbs is currently under development. This application will be available for different platforms such as Android, iOS and Windows. Making the application accessible on common devices will enable a cost-effective solution that can be easily adopted in a wide range of settings.

In the context of the rt-fMRI neurofeedback application, the use of a TCP/IP server is currently being explored. Such an option will enable the VR application to interface with the real-time analysis pipeline more robustly. Further VR features, scenarios, objects and characters will be introduced in the future, allowing, among other options, to assess the activity in different ROIs within the same scanning session. In the long run, the aim is to design experiments with complex cognitive tasks and with modulating cognitive demands that may facilitate the estimation of measures of neural capacity and efficiency in specific ROIs.

We envisage the development of a variety of other applications, using the developed multipurpose VR environment, potentially using additional devices to capture physiological changes, from electrodes for electromyography to webcams for emotion-related facial temperature changes. The VR environment is available as open-source software, under a Creative Commons license, via a GitHub repository [40].

## Funding

This work has received funding from the European Union’s Horizon 2020 research and innovation programme under the Marie Sklodowska-Curie action grant agreement No 707730.

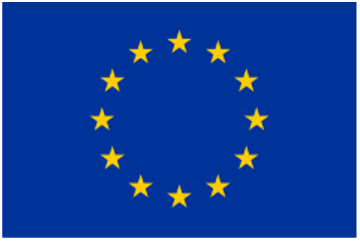

